# UNOISE2: improved error-correction for Illumina 16S and ITS amplicon sequencing

**DOI:** 10.1101/081257

**Authors:** Robert C. Edgar

## Abstract

Amplicon sequencing of tags such as 16S and ITS ribosomal RNA is a popular method for investigating microbial populations. In such experiments, sequence errors caused by PCR and sequencing are difficult to distinguish from true biological variation. I describe UNOISE2, an updated version of the UNOISE algorithm for denoising (error-correcting) Illumina amplicon reads and show that it has comparable or better accuracy than DADA2.

## Introduction

Recent examples of microbial tag sequencing experiments include the Human Microbiome Project(HMP Consortium, 2012) and a survey of the *Arabidopsis* root microbiome(Lundberg *et al.*, 2012). The experimental protocol in such studies includes amplification by PCR followed by sequencing, which introduces errors in several ways. Amplification introduces substitution and gap errors (*point errors*) due to incorrect base pairing and polymerase slippage respectively(Turnbaugh *et al.*, 2010). PCR chimeras form when an incomplete amplicon primes extension into a different biological template(Haas *et al.*, 2011). Sequencing also introduces point errors due to substitutions (incorrect base calls) and gaps (omitted or spurious base calls). Contaminants from reagents and other sources can introduce spurious species(Edgar, 2013). Spurious species can also be introduced when reads are assigned to incorrect samples due to *cross-talk*, also known as *tag switching* or *barcode switching*(Carlsen *et al.*, 2012).

The first amplicon sequencing error-correction methods were designed for pyrosequencing flowgrams(Quince *et al.*, 2011, 2009; Reeder and Knight, 2010; Rosen *et al.*, 2013). More recently, Illumina denoisers have been described including UNOISE(Edgar and Flyvbjerg, 2014), MED(Eren *et al.*, 2015) and DADA2(Callahan *et al.*, 2016). The goal of these methods is to infer accurate biological template sequences from noisy reads. This task is generally divided into two phases: 1. correcting point errors to obtain an accurate set of amplicon sequences (*denoising*) and 2. filtering of chimeric amplicons. The result is a set of predicted biological sequences that I call *ZOTUs* (zero-radius OTUs). ZOTUs are valid operational taxonomic units that are superior to conventional 97% OTUs for most purposes because they provide the maximum possible biological resolution given the data while using 97% identity may merge phenotypically different strains with distinct sequences into a single cluster(Tikhonov *et al.*, 2015; Callahan *et al.*, 2016).

The high-level strategy used by UNOISE and UNOISE2 is to cluster the unique sequences in the reads. A cluster has a centroid sequence with higher abundance plus similar sequences (*members*) having lower abundances (Fig. 1). The centroid is inferred to be correct and its members are inferred to be reads of the same template sequence containing one or more point errors. The clustering criteria in UNOISE2 have been redesigned as described below. UNOISE2 uses a one-pass clustering strategy that does not use quality (*Q*) scores and has only two parameters with pre-set values that work well on different datasets. By contrast, DADA2 uses quality scores in an iterative divisive partitioning clustering strategy based on a Poisson model with hundreds of parameters (a 4×4 transition matrix for each *Q* score) that is re-trained on each dataset. UCHIME2 and DADA2 are thus quite different, suggesting that their approaches may have complementary strengths and weaknesses. With this in mind, I will show that taking the subset of ZOTUs predicted by both algorithms reduces the number of incorrect sequences compared with using either one alone.

**Figure 1.**
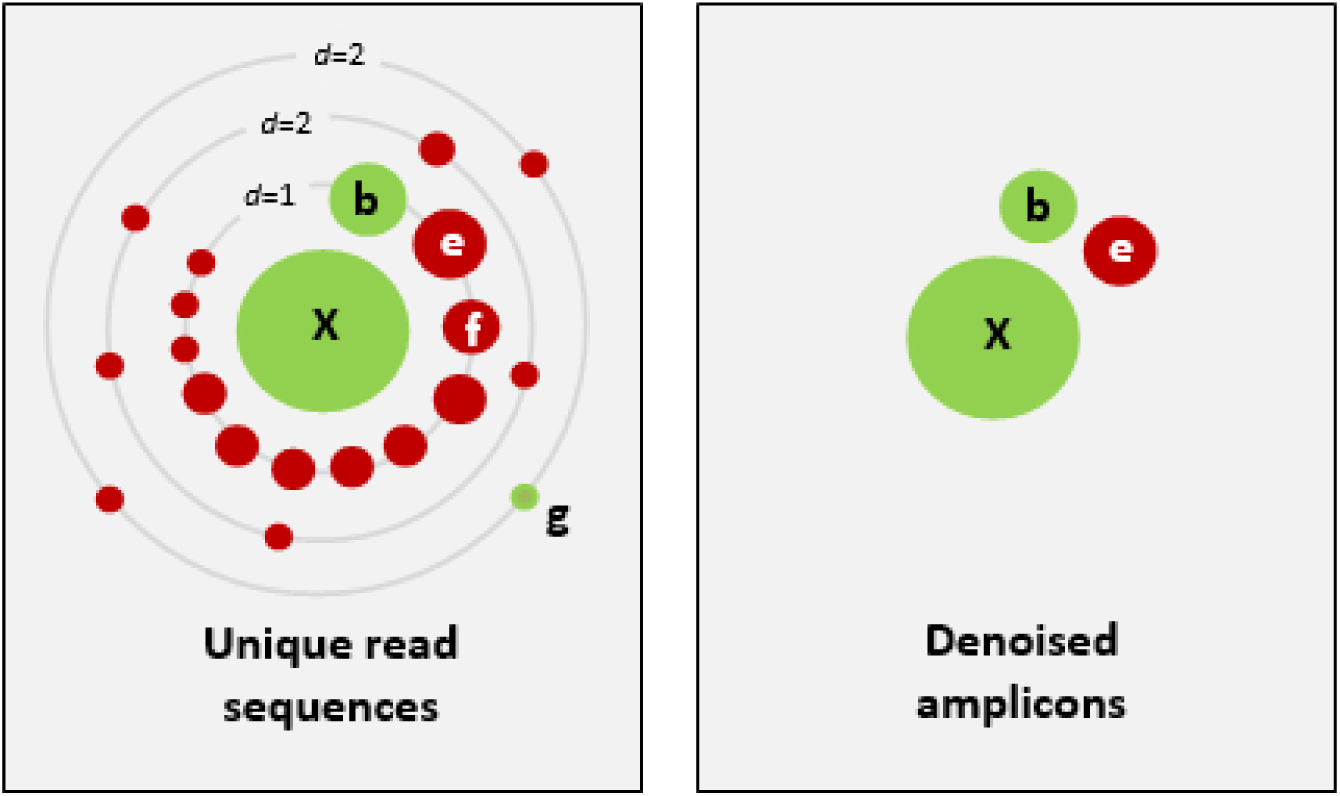
Schematic of the UNOISE2 denoising strategy. The left panel shows the neighborhood close to a high-abundance unique read sequence **X**, grouped by the number of sequence differences (*d*). Dots are unique sequences, the size of a dot indicates its abundance. Green dots are correct biological sequences; red dots have one or more errors. Neighbors with small numbers of differences and small abundance compared to **X** are predicted to be bad reads of **X.** The right panel shows the denoised amplicons. Here, **X** and **b** were correctly predicted, **e** is an error with anomalously high abundance that was wrongly predicted to be correct, **f** is an error that was correctly discarded but has an abundance almost high enough to be a false positive, and **g** is a low-abundance correct amplicon that was wrongly discarded. The abundances of **b**, **e**, and **f** are similar, illustrating the fundamental challenge in denoising: how to set an abundance threshold that distinguishes correct sequences from errors.

## UNOISE2 algorithm

Let *C* be a cluster centroid sequence with abundance *a_C_* and *M* be a member sequence of that cluster with abundance *a_M_*. Let *d* be the Levenshtein distance (number of differences including both substitutions and gaps) between *M* and *C*. The *abundance skew* of *M* with respect to *C* is defined to be *skew*(*M*, *C*)=*_aM_*/*_aC_* (Edgar *et al.*, 2011). If *M* has small enough *d* and small enough skew with respect to *C*, then it is probably an incorrect read of *C* with *d* point errors (Fig. 1). This intuition is made concrete by introducing the following function:

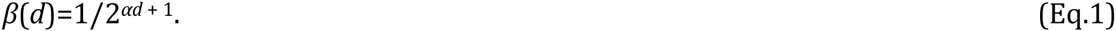

The user-settable parameter *α* is set to 2 by default, giving *β*(1)=1/8, *β*(2)=1/32, *β*(3)=1/128…. If *skew*(*M*, *C*) ≤ *β*(*d*) then *M* is a valid member of a cluster defined by *C*; i.e., *β* is the maximum skew allowed for a member with *d* differences. As *d* increases, *β* decreases exponentially, reflecting that more errors are less probable and the abundance skew should therefore be lower. The *β* function was designed by hand as a model of error abundance distributions obtained for several mock and *in vivo* Illumina datasets using the FASTX-LEARN algorithm (http://drive5.com/usearch/manual/cmd_fastx_learn.html). This is what physicists call a phenomenological model—a simple mathematical function that fits the data (and only to a very rough approximation in this case; see Discussion) without using an underlying theory.

Changing the *α* parameter trades sensitivity to small differences against an increase in the number of bad sequences which are wrongly predicted to be good. For example, setting *α*=3 gives *β*(1)=1/16, *β*(2)=1/128, *β*(3)=1/1024… which are smaller minimum skews compared to the default *α*=2. Thus, with *α*=3 a variant with *d*=1 and skew between 1/8 and 1/16 is predicted as a correct sequence while with *α*=2 it is predicted to have one error. Conversely, setting *α*=1 gives *β*(1)=1/4, *β*(2)=1/8, *β*(3)=1/16… so that a variant with *d*=1 must have a skew of at least 1/4 to be predicted as correct.

Input to the UNOISE2 algorithm is the set of unique read sequences with abundance ≥*γ*, where *γ*=4 by default. Low-abundance uniques are discarded because they are more prone to contain errors that are reproduced by chance or bias. A database of cluster centroids is initially empty. Sequences are considered in order of decreasing abundance. A sequence (*Q*) is assigned to cluster *C* if *skew*(*Q*, *C*) ≤ *β*(*d*). If no such *C* exists, *Q* becomes a new centroid. The final set of centroids are reported as the predicted amplicons. These amplicons are filtered by the UCHIME2 algorithm using denoised *de novo* mode(Edgar, 2016).

## ZOTU table construction

A table with the number of reads for each ZOTU in each sample is constructed by considering *all* reads *before* any quality filtering, including those with abundance <*γ*. The same matching criteria are used, but no new centroids are created. Thus, if a read *R* is identical to ZOTU *C*, or if *skew*(*R*, *C*) ≤ *β*(*d*), then *R* is assigned to *C*. In practice, a large majority of reads with low quality or low unique sequence abundance are due to errors in high-abundance ZOTU sequences, and this procedure thus improves sensitivity by recovering most of the reads that were discarded for the denoising step. Most of the reads not assigned to ZOTUs are usually accounted for by chimeras, which the user can verify by making a ZOTU table using predicted amplicons prior to chimera filtering.

## Sample pooling and sensitivity to rare sequences

Correct biological sequences with abundance <*γ* are lost in the denoising step and thus do not appear in the ZOTU table. I therefore recommend pooling reads from all samples in the denoising step rather than denoising each sample individually. In a typical dual-indexed sequencing run, there are ~100 samples and pooling thus increases the abundance of most correct sequences by one or two orders of magnitude, depending on how many samples contain a given strain. A sequence with abundance <4 over ~100 samples is very rare in the reads—it appears in at most three samples, in which case it would be a singleton in each, and has a maximum abundance of three in 1/100 of the samples. The data can say little about the ecological significance of this sequence. For example, it has (or should have) no significant effect on well-chosen alpha or beta diversity measures. I would therefore argue that in most cases, the loss in sensitivity due to setting *γ*=4 is inconsequential. If the user prefers to increase sensitivity at the cost of a possibly large increase in spurious ZOTUs, a smaller value of *γ* can be used.

The authors of DADA2 suggest denoising samples individually to enable detection of variants that would be lost by pooling. This happens in a scenario when a close variant (*V*) of a more dominant strain (*D*) has high abundance in one or a few samples but low abundance overall, causing *V* to be misidentified as *D* with errors. This is a valid point, and applies equally to UNOISE2. However, there are also disadvantages to this strategy. With ~100 samples, abundances are ~100× smaller in one sample and are therefore subject to much larger fluctuations which may degrade discrimination of errors from correct sequences. Some low-abundance variants may be lost that would be correctly identified by pooling, e.g., because they are singletons in some of the samples where they occur. Also, the denoiser may make different mistakes in different samples, causing a given ZOTU to contain different combinations of phenotypes. If that happens, ZOTUs are not directly comparable between samples. For example, *V* might be correctly identified as a biological variant in a few samples but misidentified as an error in others (this seems likely to occur in the motivating scenario where *V* has low overall abundance). Then, in some samples the ZOTU for *D* would contain *V* while in others *D* and *V* would be assigned to separate ZOTUs. When samples are pooled, a ZOTU will always contain the same phenotypes (hopefully, but not necessarily, just one) and this problem is avoided. With these caveats in mind, it is reasonable to try both strategies and compare the results.

## Global trimming and defining abundance

Calculating unique sequence abundance is problematic when reads of the same template sequence vary in length, e.g. because reads are truncated when the quality score drops below a threshold. Consider a case with two reads *A* and *B* where *B* is shorter but otherwise identical to *A*. Here, abundance could be defined in three different ways. (1) There are two unique sequences *A* and *B*, each with abundance one. (2) There is one unique sequence *A* with abundance two. (3) There is one unique sequence *B* with abundance two. All of these definitions have problems. With (1), a given template sequence with high abundance in the amplicons will typically have many different unique sequences with low abundances because its reads are truncated to many different lengths. With (2) the unmatched tail of *A* is considered to have the same abundance as the prefix of *A* that is identical to *B*. The tail has no support from other reads (it is effectively a singleton), but that information is lost and in practice long reads with noisy tails are assigned high abundances. With (3), the shortest sequence in a set is supported by longer sequences. This is the least bad definition: if the abundance is high, the sequence is likely to be correct. However, phylogenetically and phenotypically informative bases may be lost, and the ambiguities inherent in comparing sequences of different length must now be addressed by downstream algorithms (e.g., denoising or OTU clustering). For example, if two unique sequences differ in length by one base and have one substitution, should this count as *d*=1 (just the substitution) or *d*=2 (substitution plus terminal gap)? If large variations in length are allowed, then the phylogenetic and phenotypic resolution of the sequences may vary substantially, degrading the comparability of ZOTUs or OTUs to each other for calculating diversity, predicting taxonomy and so on. These problems are avoided by ensuring that reads of the same template sequence have the same length (*global trimming*, implying that reads of the same template should be globally alignable, though more distantly related sequences need not be). The simplest method for global trimming is to truncate all reads to the same length. This is not usually necessary with overlapping Illumina paired-end reads that have been merged by a paired-read assembler. In this case, the merged sequence always terminates at the reverse primer which guarantees that reads of the same template will have the same length regardless of variations in amplicon length between different species. If multiple primers were used which do not bind to the same locus, then trimming is required to ensure that reads of the same template amplified by different primers start and end at the same position in the biological sequence. Primer-binding bases should be discarded from the reads because PCR tends to induce substitutions at mismatched positions; in most cases this is easily accomplished by discarding a fixed number of bases (the primer lengths) from each end of the sequence. There is no need to explicitly match the primer sequence in order to trim it unless there are multiple primers binding to different loci.

## Quality filtering and paired read merging

UNOISE2 *per se* does not use quality scores. However, denoising is more effective with quality-filtered reads because sequencing error bias can cause some errors to have sufficiently high abundances that they could be mistaken for biological variants, and these often have lower quality scores. I therefore recommend discarding reads with ≥1 expected errors so that the most probable number of errors is zero for all reads(Edgar and Flyvbjerg, 2014). Paired reads should be merged using a Bayesian assembler before expected error filtering to exploit the improved base calls and posterior error probabilities obtained in the overlapping region(Edgar and Flyvbjerg, 2014). Methods that truncate at varying lengths based on quality scores, e.g. the QIIME quality filter described in(Bokulich *et al.*, 2013), should not be used (see Global trimming above).

## Validation

I compared UNOISE2 and DADA2 on three mock and three *in vivo* datasets (Table 1). Two of the mock datasets use a community designed(Haas *et al.*, 2011) for the Human Microbiome Project and one is the Extreme dataset(Callahan *et al.*, 2016) used to validate DADA2. DADA2 was not tested on the ITS data as it is not designed for amplicons with larger variations in length. On the vagina reads, DADA2 predicted 187 amplicon sequences twice, differing only in length. I discarded the shorter version of each sequence.

**Table 1.**
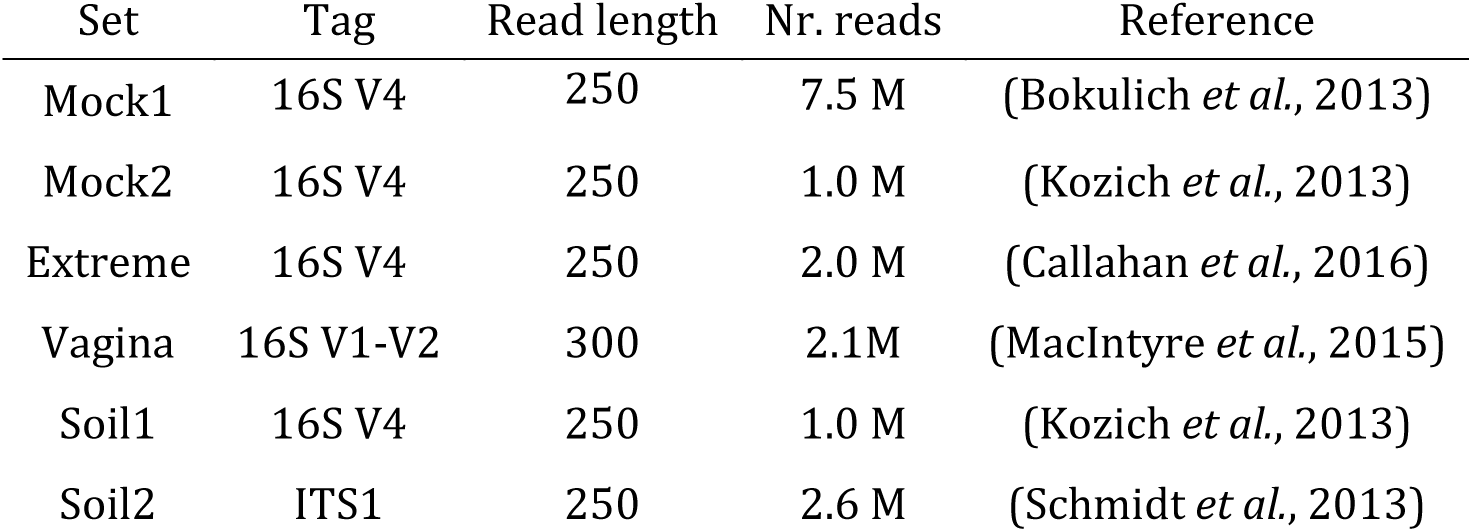
Datasets used for testing. All datasets contain MiSeq paired-end reads. A random subset of 1M read pairs was extracted from the Mock2 and Soil1 datasets because DADA2 failed to converge in the model estimation step when all reads were input. Mock1 and Mock2 are the mock communities with 21 strains(Haas *et al.*, 2011) used to validate sequencing protocols in the Human Microbiome Project. The Extreme mock community has 27 strains(Callahan *et al.*, 2016).

A ZOTU was classified as follows. If it is 100% identical to a known sequence in the designed mock community, it is *Perfect*. If it is <100% and ≥97% to a known mock sequence, it is *Noisy*. If it is 100% identical to a sequence in the SILVA(Pruesse *et al.*, 2007) 16S database v123 or the UNITE(Kõljalg *et al.*, 2013) ITS database v04.11.2015, it is *Exact*, if >97% identical *Good*, and if none of the above then *Other*. On mock data, the mock reference database is considered before the large database (SILVA or UNITE).

A *Noisy* ZOTU is most likely to have uncorrected errors, though it could be a valid biological variant that is present in the community but missing from the mock reference database, e.g. due to an unknown paralog or an impure cell line used to make the mock sample. An *Exact* ZOTU is probably correct, though it could be an error that reproduces a sequence found in the database (this is more likely that it might appear as explained in the discussion of chimera filtering below). An *Exact* ZOTU in a mock sample is <97% with all known mock sequences and 100% identical to a SILVA sequence and is therefore very likely to be a correct biological sequence due to contamination or cross-talk. A ZOTU classified as *Good* or *Other* could be a correct sequence that is missing from SILVA or UNITE, or it could have errors. N1, N2 and N3 are the number of ZOTUs for which the nearest ZOTU has 1, 2 and 3 differences, respectively. These indicate how many close variants were predicted, but are difficult to interpret on *in vivo* samples because those variants could be true biological sequences or uncorrected errors. OTUs were constructed by clustering ZOTUs with UCLUST(Edgar, 2010) at 97% identity. The average number of ZOTUs per OTU gives an indication of the increased resolution achieved at 100% identity.

Results are summarized in Table 2. On the Mock1 dataset, the UNOISE2 results are better. Almost half (22/50) of the DADA2 ZOTUs are *Noisy* while UNOISE2 predicted 23/29 *Perfect* amplicons, 3/29 *Exact* and no *Noisy*. On the other mock datasets, the programs have similar performance. On Mock2, most ZOTUs are *Perfect* or *Exact*. Both predict ~100 *Exact* amplicons which are explained by high-frequency MiSeq cross-talk (manuscript in preparation). Taking the subset of ZOTUs predicted by both programs (U&D) retains most of the *Perfect* and *Exact* amplicons in all datasets, while discarding most of the *Noisy* amplicons and many of the *Good* and *Others*. This supports the hypothesis that *Good* and *Other* probably contain a mix of errors and correct sequences missing from the reference database. The number of *d*=1 neighbors (N1) is substantially higher for UNOISE2 in Soil1. This is mostly accounted for by the larger numbers of chimeras predicted by DADA2, many of which have only a single difference with their closest putative parent. If these DADA2 chimera predictions are mostly false positives, as I argue under Chimera filtering below, then UNOISE2+UCHIME2 has better sensitivity to close variants. However, this cannot be established with certainty and it is also possible that UCHIME2 failed to filter a substantial number of valid chimeras, in which case it would be better to use non-default parameters for UCHIME2 specifying equivalent criteria to DADA2.

**Table 2.** Results on test datasets. See main text for explanation of column headings and discussion of the results. U&D is the consensus of UNOISE2 and DADA2, i.e. ZOTUs predicted by both algorithms.

| Set | Method | ZOTUs / OTUs | Perfect | Noisy | Exact | Good | Other | N1 | N2 | N3 |
| --- | --- | --- | --- | --- | --- | --- | --- | --- | --- | --- |
| Mock1 | UNOISE2 | 29 / 27 (1.1) | 22 | 0 | 5 | 2 | 0 | 2 | 1 | 0 |
|  | DADA2 | 50 / 25 (2.0) | 23 | 22 | 3 | 1 | 1 | 28 | 0 | 3 |
|  | U&D | 26 / 24 (1.1) | 22 | 0 | 2 | 1 | 1 | 3 | 0 | 0 |
| Mock2 | UNOISE2 | 126 / 98 (1.3) | 21 | 1 | 98 | 7 | 0 | 17 | 8 | 8 |
|  | DADA2 | 134 / 121 (1.1) | 22 | 3 | 102 | 3 | 4 | 6 | 2 | 6 |
|  | U&D | 105 / 95 (1.1) | 21 | 1 | 80 | 1 | 0 | 3 | 2 | 4 |
| Extreme | UNOISE2 | 31 / 22 (1.4) | 24 | 2 | 5 | 0 | 0 | 9 | 2 | 0 |
|  | DADA2 | 26 / 20 (1.3) | 24 | 0 | 2 | 0 | 0 | 5 | 2 | 0 |
|  | U&D | 25 / 19 (1.3) | 23 | 0 | 2 | 0 | 0 | 5 | 2 | 0 |
| Vagina | UNOISE2 | 464 / 224 (2.1) | - | - | 251 | 213 | 18 | 101 | 82 | 66 |
|  | DADA2 | 432 / 263 (1.6) | - | - | 268 | 122 | 42 | 78 | 59 | 41 |
|  | U&D | 309 / 185 (1.7) | - | - | 219 | 71 | 19 | 48 | 52 | 36 |
| Soil1 | UNOISE2 | 10286 / 4767 (2.2) | - | - | 2624 | 5737 | 2313 | 1379 | 2046 | 1933 |
|  | DADA2 | 7531 / 4824 (1.6) | - | - | 2073 | 2350 | 3108 | 488 | 925 | 1020 |
|  | U&D | 7148 / 4508 (1.6) | - | - | 2015 | 2280 | 1097 | 434 | 906 | 1010 |
| Soil2 | UNOISE | 1369 / 540 (2.5) | - | - | 107 | 289 | 950 | 755 | 242 | 130 |

Results on the Extreme mock community are summarized in Table 3. Shaded rows are ZOTUs which are not found in the mock reference database but are exact matches to SILVA; these are probably contaminants not in the designed community or variants missing from the reference database. All five of these were identified by UNOISE2, two of which were also found by DADA2. Two of the designed strains were predicted to have one *Perfect* ZOTU and one *Noisy* ZOTU by UNOISE2, otherwise all ZOTUs for designed strains were *Perfect* from both programs. UNOISE2 failed to identify one low-abundance designed strain, *P. copri*, which was successfully identified by DADA2.

**Table 3.**
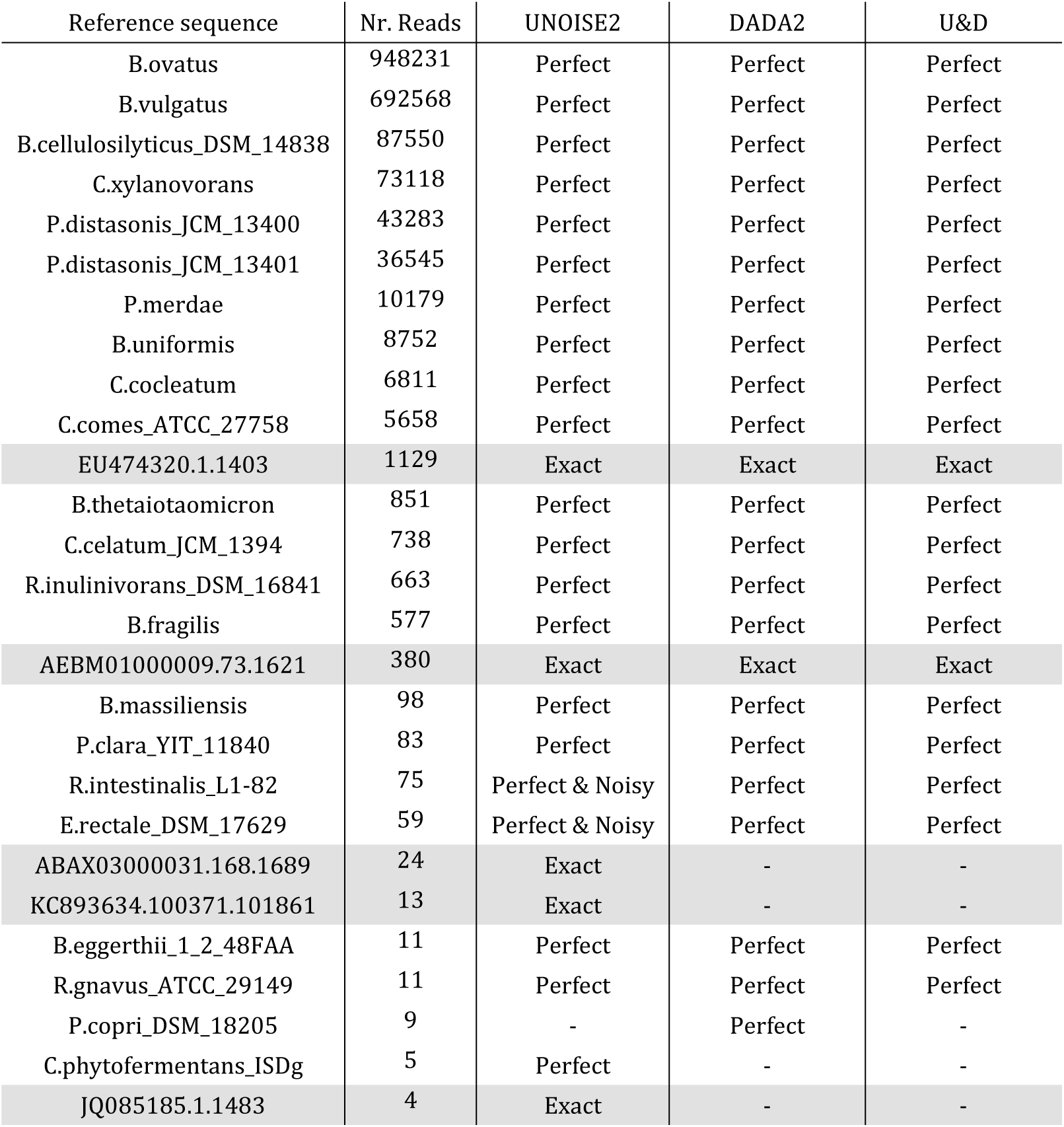
Results on the Extreme mock community. The table shows strains identified by UNOISE2 or DADA2. Shaded rows are ZOTUs which are not found in the mock reference database but are exact matches to SILVA. U&D is ZOTUs predicted by both UNOISE2 and DADA2.

| Reference sequence | Nr. Reads | UNOISE2 | DADA2 | U&D |
| --- | --- | --- | --- | --- |
| B.ovatus | 948231 | Perfect | Perfect | Perfect |
| B.vulgatus | 692568 | Perfect | Perfect | Perfect |
| B.cellulosilyticus_DSM_14838 | 87550 | Perfect | Perfect | Perfect |
| C.xylanovorans | 73118 | Perfect | Perfect | Perfect |
| P.distasonis_JCM_13400 | 43283 | Perfect | Perfect | Perfect |
| P.distasonis_JCM_13401 | 36545 | Perfect | Perfect | Perfect |
| P.merdae | 10179 | Perfect | Perfect | Perfect |
| B.uniformis | 8752 | Perfect | Perfect | Perfect |
| C.cocleatum | 6811 | Perfect | Perfect | Perfect |
| C.comes_ATCC_27758 | 5658 | Perfect | Perfect | Perfect |
| EU474320.1.1403 | 1129 | Exact | Exact | Exact |
| B.thetaiotaomicron | 851 | Perfect | Perfect | Perfect |
| C.celatum_JCM_1394 | 738 | Perfect | Perfect | Perfect |
| R.inulinivorans_DSM_16841 | 663 | Perfect | Perfect | Perfect |
| B.fragilis | 577 | Perfect | Perfect | Perfect |
| AEBM01000009.73.1621 | 380 | Exact | Exact | Exact |
| B.massiliensis | 98 | Perfect | Perfect | Perfect |
| P.clara_YIT_11840 | 83 | Perfect | Perfect | Perfect |
| R.intestinalis_L1-82 | 75 | Perfect & Noisy | Perfect | Perfect |
| E.rectale_DSM_17629 | 59 | Perfect & Noisy | Perfect | Perfect |
| ABAX03000031.168.1689 | 24 | Exact | - | - |
| KC893634.100371.101861 | 13 | Exact | - | - |
| B.eggerthii_1_2_48FAA | 11 | Perfect | Perfect | Perfect |
| R.gnavus_ATCC_29149 | 11 | Perfect | Perfect | Perfect |
| P.copri_DSM_18205 | 9 | - | Perfect | - |
| C.phytofermentans_ISDg | 5 | Perfect | - | - |
| JQ085185.1.1483 | 4 | Exact | - | - |

## Chimera filtering

Validation of chimera filtering for ZOTUs raises issues that can be neglected with 97% OTUs. For example, the most abundant unique sequence in Mock1 occurs in 2.8×10^6^ reads. This sequence is from *Staphylococcus epidermidis* and *S. aureus*, which are both present in the community and have identical V4 sequences. The sequence length is 233nt after merging of read pairs and trimming the primer-binding bases. There are 3×233=699 possible variants with one substitution, all of which are present in the reads. Several of these variants can be reconstructed exactly as two-segment chimeras from known sequences in the mock community, and these potentially chimeric reads therefore cannot be distinguished from *d*=1 point errors. Presumably, in some cases the same unique sequence is generated by point errors (which are likely to be reproduced several times with such a large read depth) and by one or more chimera formation events (chimeras formed by different parents or different cross-over points can have identical sequences). The point errors could be due to PCR or to sequencing; most likely both. In such cases, the unique sequence cannot be exclusively classified as a chimera or as a point error because it is found in both types of read. Also, the reference database for the HMP mock community (21 strains) has 115 different 16S sequences, an average of 5.5 distinct 16S sequences per strain. While having multiple 16S operons per genome is common, 16S paralogs usually have the same sequence(Acinas *et al.*, 2004) and it therefore seems unlikely that all of the sequences in the reference database are present in the genomes of the ATCC cell lines specified for the community (this is hard to verify because it is not documented how the database was made, to the best of my knowledge—it is also possible that there are variants in the database found by Sanger sequencing and explained by impure cell lines). Extraneous sequences are problematic for validation because they increase the likelihood of incorrect inferences of chimeras and can cause false inferences that amplicon sequences are non-chimeric and correct. For example, if there is an *S. aureus* variant with one difference in the reference database, it will be present in the reads (because *all d*=1 variants are present), even if that variant is not present in the sample.

UCHIME2 and the DADA2 removeBimeraDenovo function use similar algorithms for chimera detection in denoised amplicons, but there are differences in the details which are important in practice. Both construct a model of the query sequence from segments of two other amplicons (the *parents*). UCHIME2 requires that the model is identical to the query sequence while DADA2 allows one difference with the model if the query has four or more differences with both parents. Also, UCHIME2 requires an abundance ratio of at least two; i.e. the least abundant parent used to make the model must be at least twice as abundant as the query (because parents undergo at least one more round of amplification than the chimera), while DADA2 requires only that the parents are more abundant than the query. DADA2 will therefore tend to discard more sequences: those having one difference in the model or an abundance ratio between one and two.

Comparing DADA2 and UCHIME2 chimera filtering on mock data would ideally use an independent method to classify sequences as non-chimeric or chimeric by comparison with the mock sequence reference database. However, as discussed above, it is not possible in general to determine whether a sequence is chimeric and / or has point errors, especially when the number of differences is small. A sequence cannot be reliably classified as non-chimeric unless it is identical to a reference sequence(Edgar, 2016), and amplicons with uncorrected point errors therefore cannot be reliably classified. These issues should not be circumvented by choosing to consider only chimeras with more differences (which can be distinguished from point errors more reliably), because low-divergence chimeras are the most common(Edgar, 2016) and are therefore the most important in the context of denoising, which by definition attempts to resolve low-divergence biological differences. Given these considerations, it is clear that the (debatably) independent reference-based methods which could be considered, i.e. UCHIME2 in denoised reference mode, UPARSE-REF(Edgar, 2013) or the seqs.error command in mothur(Schloss *et al.*, 2009), are no more or less trustworthy than the methods being tested despite the apparent advantage of using the database, and would likely be biased in favor of one of the algorithms, depending on which method and parameters were chosen (e.g., gap and cross-over penalties for scoring chimeric alignments). Note also that a mock reference database could contain errors and is almost certainly incomplete due to contaminants and cross-talk in the reads, and it is therefore plausible that *de novo* methods, which can detect unexpected sequences, could construct a more accurate database than the "official" reference. Simulated data would have similar issues with the additional problem that the simulation might not be sufficiently realistic (note the circular problem of learning parameters for such a model from real data when chimeras and point errors due to sequencing and PCR cannot be distinguished). In conclusion, robustly measuring chimera filtering accuracy of denoisers using a gold standard is not possible because there is no method for identifying or simulating chimeras that is demonstrably better than the algorithms to be tested.

I compared the results of DADA2 and UCHIME2 to each other by running both removeBimeraDenovo and UCHIME2 on amplicons predicted by DADA2. On the low-diversity mock community data, the algorithms were in agreement: both filtered the same sequences. However, there were substantial differences on the *in vivo* data, most obviously on the high-diversity 16S soil sample (Fig. 2). Here, DADA2 reported 939 of 8,470 amplicons as chimeric while UCHIME2 filtered only 401, less than half as many. Of these 401, all but one were also filtered by DADA2. Of the 539 filtered only by DADA2, 282 had abundance ratios between one and two and the remaining 257 had one difference (Fig. 2). The 539 sequences filtered only by DADA2 must be false positives by DADA2 or false negatives by UCHIME2. I believe that a large majority of these are probably false positives by DADA2, and that most likely many of the 400 filtered by both are also false positives, while acknowledging that my reasoning makes assumptions and approximations that are difficult to test.

**Figure 2.**
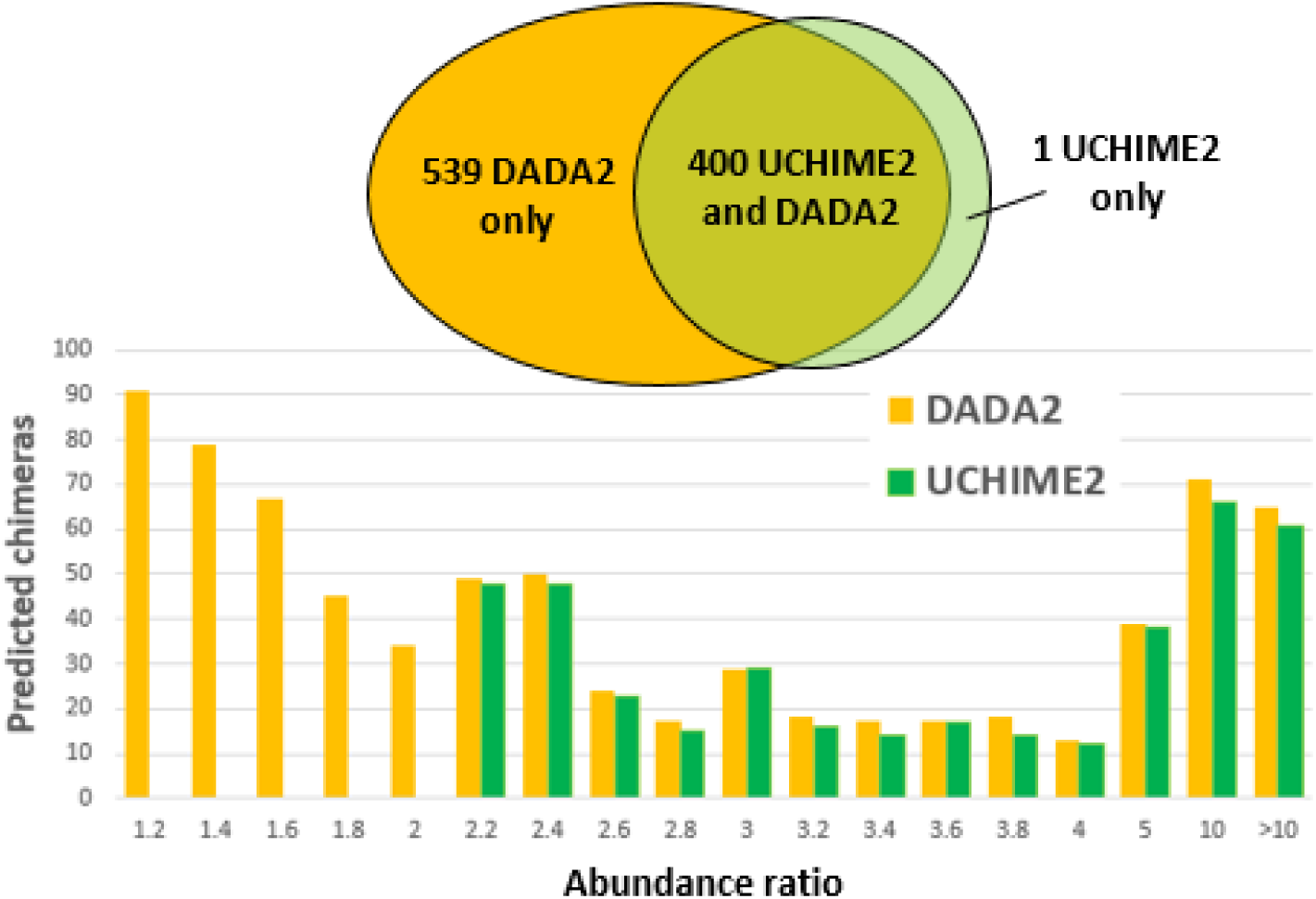
Chimeras predicted by DADA2 and UNOISE2 on the Soil1 dataset. Each histogram bar gives the number of predicted chimeras in an abundance ratio (AR) range labeled by its upper value, so the first bin contains chimeras with 1.0≤AR<1.2, the second 1.2≤AR<1.4 and so on. The last bin has all chimeras with AR>10. Notice that DADA2 predicts more than twice as many chimeras as UCHIME2, many of which have AR<2, while most chimeras would be expected to have AR≥2 because the parents undergo at least one more round of PCR amplification. In the bins with AR≥2 the two programs agree on most predictions. DADA2 predicts a few more in each bin because it sometimes allows one difference in the chimeric model built from the putative parent sequences while UCHIME2 always requires an exact match.

The 257 cases where the model has one difference necessarily imply incorrect predictions by DADA2—if the query is chimeric then one of the three amplicons (query plus parents) has an uncorrected point error causing the difference, and if the query is not in fact chimeric then it is a false positive. (For a loophole in this argument, see What are amplicons? below). I believe that false positive chimeras will have a much higher frequency than uncorrected point errors, given the high accuracy of DADA2 on most of the mock datasets and the observation that fake chimeric models are very common, especially when differences are allowed(Edgar, 2016).

Most of the 282 sequences filtered by DADA2 with abundance ratios between one and two are probably also false positives. The argument here is based on the abundance ratio distribution expected for true chimeras. Suppose there are *N* rounds of PCR. A chimera created in the *k*th round should have an abundance ratio of ~2^*k*^ because the parents are duplicated by all *N* rounds while the chimera is duplicated by the *N*–*k* remaining rounds after it is formed. The lowest observed abundance ratios should therefore be ~2 due mostly to chimeras formed in the first round. The frequency of ratios between two and one should fall as the ratio decreases, and ratios close to one should have very low frequency. If the rate of chimera formation is constant in each round, then ~1/*N* of the chimeras will form in the first round. If fluctuations in the abundance ratio are equally likely to give values <2 and >2, then approximately half of the chimeras formed in the first round will have abundance ratio <2, i.e. 1/(2*N*). (E.g., to a first approximation the distribution could be normal with a mean of 2). The soil sample was amplified using 30 PCR cycles, so this rough estimate predicts that ~1/60th (2%) of chimeras will have abundance ratios <2, with a small minority of these close to one. I suspect that chimeras are much more likely to form in later rounds because there are more amplicons, in which case this estimate is conservative. Either way, 2% is a reasonable estimate for the upper bound on the number of chimeras we should expect with abundance ratios <2. These expectations are very different from the DADA2 predictions (Fig. 2). A majority (282/539=52%) have abundance ratio <2., and 90/539=17% have abundance ratios <1.2. The frequency increases rather than decreases as the abundance ratio drops from two to one. These observations are difficult to reconcile with true chimeras, but are readily explained if most of the predictions are false positives due to fake models.

If many or most of the DADA2 predictions with abundance ratio <2 are false positives, then many of the predictions with small ratios >2, say in the range two to three, are probably also false positives because the false positive rate should be approximately independent of the abundance ratio when the ratio is small (with larger ratios, the number of candidate parents is reduced, which may reduce the rate of false positives because there are fewer ways to make fakes). Given that UCHIME2 agrees with 400/657 of the DADA2 chimera predictions with ratios >2, it seems likely that many of the UCHIME2 predictions are also false positives, despite using more stringent parameters (no differences allowed in the model, abundance ratio ≥2).

## What is an amplicon?

To this point, I have described denoising as prediction of amplicon sequences followed by chimera filtering. This glosses over a complication that is important for chimera identification. If PCR point errors are often found in the reads, then we want to correct them in order to recover the biological sequences. But for optimal chimera identification at single-base resolution we need all of the amplicon sequences generated in the PCR reaction, including those with point errors because we want to find chimeras whose parent segments have PCR point errors. Ideally, a denoiser pipeline would therefore 1. correct sequencer error, leaving PCR point errors; 2. filter chimeras; then 3. correct PCR point errors. However, I do not believe that this is a tractable approach because point errors due to sequencing generally cannot be distinguished from point errors due to PCR. So in practice, denoisers implicitly attempt to remove *all* point errors before chimera filtering, which might degrade chimera filtering. If you believe that PCR point errors are common, then it might be reasonable to allow one or even more differences between the chimera and its model. This is a possible justification for the DADA2 chimera identification criteria, though no motivation for allowing a difference has been given by the authors, to the best of my knowledge. However, this design is likely to increase the number of false-positive chimera predictions. I believe that it is probably better to assume that chimeras with PCR point errors are rare and those that do form are more likely to be created in later rounds and most will be discarded because they have very low abundances <*γ*, in which case it is better to require an exact match to the model. Hence the choice of default parameters in UCHIME2.

## Discussion

Denoising exploits the observation that a low-abundance sequence that is very similar to a high-abundance sequence is likely to be an error. The fundamental challenge of denoising is determining an abundance threshold that distinguishes a correct sequence from an error. Error frequencies vary due to biases which cannot be accurately predicted and to fluctuations due to sampling effects which are predictably present but have unpredictable values for any given sequence. Outliers are common, explained by fluctuations, PCR point errors and non-random sequencing error; see e.g. (Schirmer *et al.*, 2015). Non-random sequencing error cannot be explained by measurable biases such as tendencies to make certain base call substitution errors. PCR point errors are amplified in subsequent rounds of PCR, so while it is plausible that point errors are generated by an approximately Poisson process, they are then exponentially amplified causing anomalously high abundances by factors of ~2^*k*^ for *k* =1 … *N*-1 if there are *N* rounds. These observations necessitate setting very conservative thresholds which fail to identify low-abundance biological variants in order to suppress anomalously high-frequency errors which are very common in practice. In other words, denoisers must lean strongly towards “lumping” rather than “splitting” ZOTUs. In the case of DADA2, this is reflected in the astronomically small default value of its OMEGA_A parameter (10^−20^). Abundance *p*-values above OMEGA_A are considered to indicate an error in the sequence, so *p*=10^−19^ is bad but *p*=10^−21^ is good. In the case of UNOISE2, this is reflected in the parameters of the *β* function which were chosen to suppress outliers rather than typical errors. With 16S and ITS data, this approach gives largely accurate and useful results despite its unavoidable limitations because close variants are relatively rare and merging them is relatively benign. It is clearly better to accept that some *d*=1 or *d*=2 variants will be lumped into the same ZOTU than to use traditional clustering at 97% identity, which always lumps variants with *d*≤7 into the same OTU when the sequence length is ~250nt.

